## Supplemental Table and Figures for "T-follicular helper cells are epigenetically poised to transdifferentiate into T-regulatory type-1 cells"

**Supplementary Table 1**. DE expression of TR1/TFH/Treg-relevant genes in pMHCII-NP-induced Tet+ vs. vaccine-induced TFH cells (111)

| Gene type | Genes (protein) | Described in TR1 (ref) | Described in other Treg types (ref) | Described in TFH (ref) | DE in Tet+ (vs. TFH) |
|---|---|---|---|---|---|
| Cell adhesion molecules | Cd226 | + (1) | ND | ND | +++ |
| | Itga2 (CD49b) | + (1) | ND | ND | ++ |
| | Itgae (CD103) | + (2) | + (3) | ND | +++ |
| | Ocln | ND | ND | ND | +++ |
| | S1pr2 | ND | ND | + (4) | - |
| | Sell (CD62L) | - (5) | + (6) | - (7) | +++ |
| | Selplg (Psgl1) | ND | ND | - (8) | +++ |
| Chemokyne receptors | Ccr5 | + (9) | + (10) | + (11) | +++ |
| | Ccr7 | ND | + (12) | - (13) | +++ |
| | Cxcr3 | + (14) | + (12) | In some subsets (15) | +++ |
| | Cxcr4 | ND | ND | + (16) | NS |
| | Cxcr5 | ND | ND | + (17) | - - |
| Co-estimulatory molecules | Cd28 | + (18) | ND | + (19) | + |
| | Cd40lg | ND | ND | + (20) | + |
| | Icos | + (21) | + (22) | + (23) | - |
| | Klrk1 (NKG2D) | ND | + (24) | ND | NS |
| | Sh2d1a (SAP) | ND | ND | + (25) | - - |
| | Tnfrsf4 (Ox40) | + (26) | + (27) | + (28) | ++ |
| | Tnfrsf18 (GITR) | + (18) | + (29) | ND | + |
| | Tnfsf4 (Ox40L) | ND | ND | ND | +++ |
| Co-inhibitory molecules | Ctla4 | + (30) | + (31) | + (32) | ++ |
| | Fasl | ND | + (33) | ND | ++ |
| | Havcr2 (TIM-3) | + (34) | + (35) | + (36) | +++ |
| | Lag3 | + (1) | + (37) | ND | +++ |
| | Pdcd1 (PD-1) | + (9) | + (38) | + (17) | - - |
| | Tigit | + (39) | + (40) | + (41) | NS |
| Cytokines | Ebi3 (IL27b) | ND | + (42) | ND | + |
| | Ifng | + (43) | ND | ND | +++ |
| | Il10 | + (44) | + (45) | + (46) | +++ |
| | Il21 | + (47) | ND | + (48) | NS |
| | Il4 | - (44) | ND | + (49) | - |
| | Il5 | + (44) | ND | ND | NS |
| | Mcub (Areg) | + | + (50) | ND | NS |
| | Tgfb1 | + (44) | + (45) | ND | NS |
| Cytokine receptors | Il10ra | + (51) | + (52) | ND | ++ |
| | Il12rb2 | ND | + (53) | ND | - - |
| | Il21r | + (47) | + (54) | + (48) | - |
| | Il27ra | + (55) | + (56) | + (57) | + |
| | IL7r (CD127) | - (21) | - (58) | - (59) | +++ |
| | IL17rc | ND | ND | ND | NS |
| | Il2ra (CD25) | - (18) | + (60) | - (61) | +++ |
| | Tgfbr1 | - (51) | + (62) | ND | - |
| | Tgfbr2 | - (51) | + (63) | ND | NS |
| | Tgfbr3 | - (51) | ND | ND | ++ |
| | Atf6 | - (51) | ND | ND | NS |
| | Ahr | + (64) | ND | ND | +++ |
| | Ajuba | ND | ND | ND | +++ |
| | Ascl2 | ND | ND | + (65) | - |
| | Bach2 | - (66) | + (67) | - (68) | ++ |
| | Batf | + (69) | ND | + (70) | - |
| | Bcl6 | ND | ND | + (71) | - - |
| | Bhlhe40 | + (51) | ND | ND | ++ |
| | Bmyc | - (51) | ND | ND | NS |
| | Cbfa2t3 | ND | ND | ND | - |
| | Cebpa | ND | ND | + (72) | - - |

| | | | | | |
|---|---|---|---|---|---|
| Transcription factors | Dbp | - (51) | ND | ND | NS |
| | E2f1 | + (51) | + (73) | ND | NS |
| | Egr2 | + (74) | ND | ND | - |
| | Elk4 | - (51) | ND | ND | - |
| | Eomes | + (75) | + (76) | ND | ++ |
| | FoxP1 | ND | ND | - (77) | + |
| | FoxP3 | ND | + (78) | ND | - - |
| | Grhl1 | ND | ND | ND | NS |
| | Hmgb2 | + (51) | ND | ND | NS |
| | Id2 | + (51) | + (79) | - (80) | ++ |
| | Id3 | - (51) | + (79) | + (80) | - |
| | Irf1 | + (69) | ND | ND | + |
| | Irf4 | + (81) | + (3) | + (82) | ++ |
| | Jdp2 | ND | ND | ND | - - |
| | Klf2 | ND | ND | - (83) | +++ |
| | Lef1 | ND | ND | + (84) | NS |
| | Lilrb4a | ND | + (85) | ND | NS |
| | Maf | + (47) | ND | + (86) | - |
| | Myb | - (51) | + (87) | ND | NS |
| | Mybl2 | + (51) | ND | ND | - - |
| | Myc | - (51) | - (88) | ND | NS |
| | Nfia | ND | ND | ND | - - |
| | Nfil3 | + (34) | ND | ND | ++ |
| | Nr1h3 (LXR$\alpha$) | + (51) | + (89) | ND | - |
| | Pax5 | ND | ND | ND | - - |
| | Pax9 | ND | ND | ND | ++ |
| | Pou2af1 (OcaB) | ND | ND | ND | - |
| | Prdm1 (Blimp-1) | + (90) | ND | + (91) | +++ |
| | Rbpj | + (51) | + (92) | ND | + |
| | Runx2 | + (51) | ND | ND | ++ |
| | Rora | + (51) | ND | - (93) | - - |
| | S1pr1 | ND | ND | - (83) | ++ |
| | Six5 | ND | ND | ND | NS |
| | Sox4 | - (51) | - (94) | ND | NS |
| | Sox8 | ND | ND | ND | - - |
| | Stat1 | + (95) | ND | + (96) | NS |
| | Stat3 | + (97) | ND | + (23) | NS |
| | Stat4 | ND | - (98) | + (99) | - |
| | Tbx21 (T-bet) | + (75) | ND | - (84) | ++ |
| | Tcf7 | ND | ND | + (84) | NS |
| | Tox2 | ND | ND | + (80) | - - |
| | Vdr | ND | + (100) | ND | NS |
| | Zbtb16 (PLZF) | + (51) | ND | ND | NS |
| Secretion proteins | Chgb | ND | ND | + (101) | NS |
| | Gzmb (Granzyme B) | + (102) | + (103) | ND | +++ |
| Enzymes | Cblb | ND | + (104) | ND | + |
| | Entpd1 (CD39) | + (105) | + (106) | ND | +++ |
| | Itk | + (107) | + (108) | ND | - |
| | Nt5e (CD73) | + (105) | + (106) | + (109) | NS |
| | Serpinb6b | + (51) | + (110) | ND | NS |
| | Serpinb9 | + (51) | ND | ND | NS |

| | |
|---|---|
| +++ | FC>4 |
| ++ | 4>FC>2 |
| + | 2>FC>1 |
| NS | 1>FC>-1 |
| - | -1>FC>-2 |
| - - | -2>FC>-4 |
| - - - | FC<-4 |
| ND | Not determined |

Suppl. Fig. 1

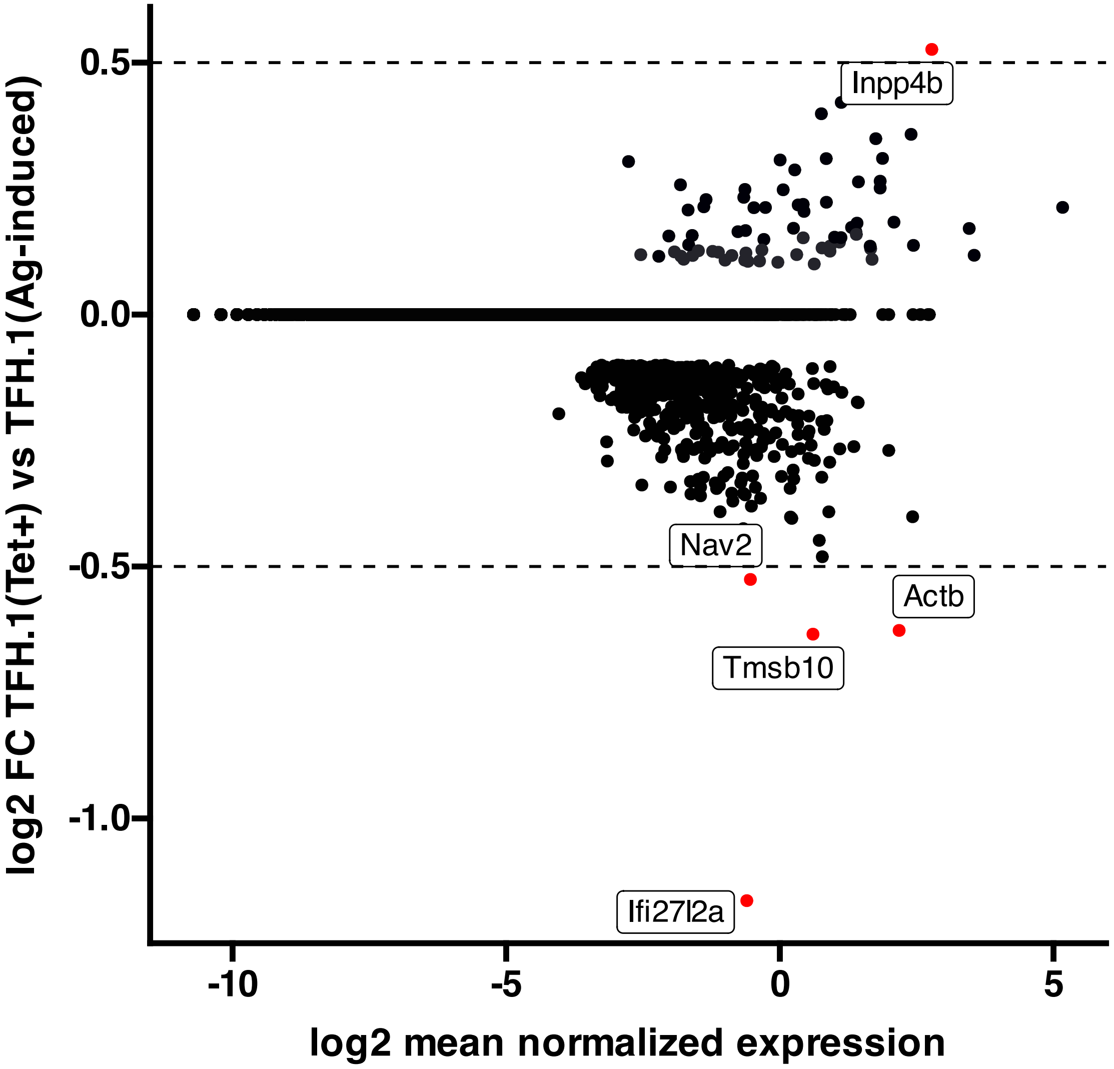

log2 FC TFH.1(Tet+) vs TFH.1(Ag-induced)
0.5
0.0
-0.5
-1.0
-10
-5
0
5
log2 mean normalized expression
Inpp4b
Nav2
Actb
Tmsb10
Ifi27l2a

**Supplementary Figure 1. High transcriptional similarity between pMHCII-NP- and vaccine-induced TFH.1 cells.** Figure corresponds to an MA plot comparing the transcriptome of the TFH.1 cells from the KLH-induced TFH cell pool and the TFH-like cells contained within the BDC2.5mi/I-$A^{g7}$-NP-induced Tet+ pool. There were only 5 differentially expressed genes (|log2FC| > 0.5 and adjusted P<0.05; *Actb, Ifi27l2a, Inpp4b, Nav2* and *Tmsb10*) between these two subsets.

Suppl. Fig. 2

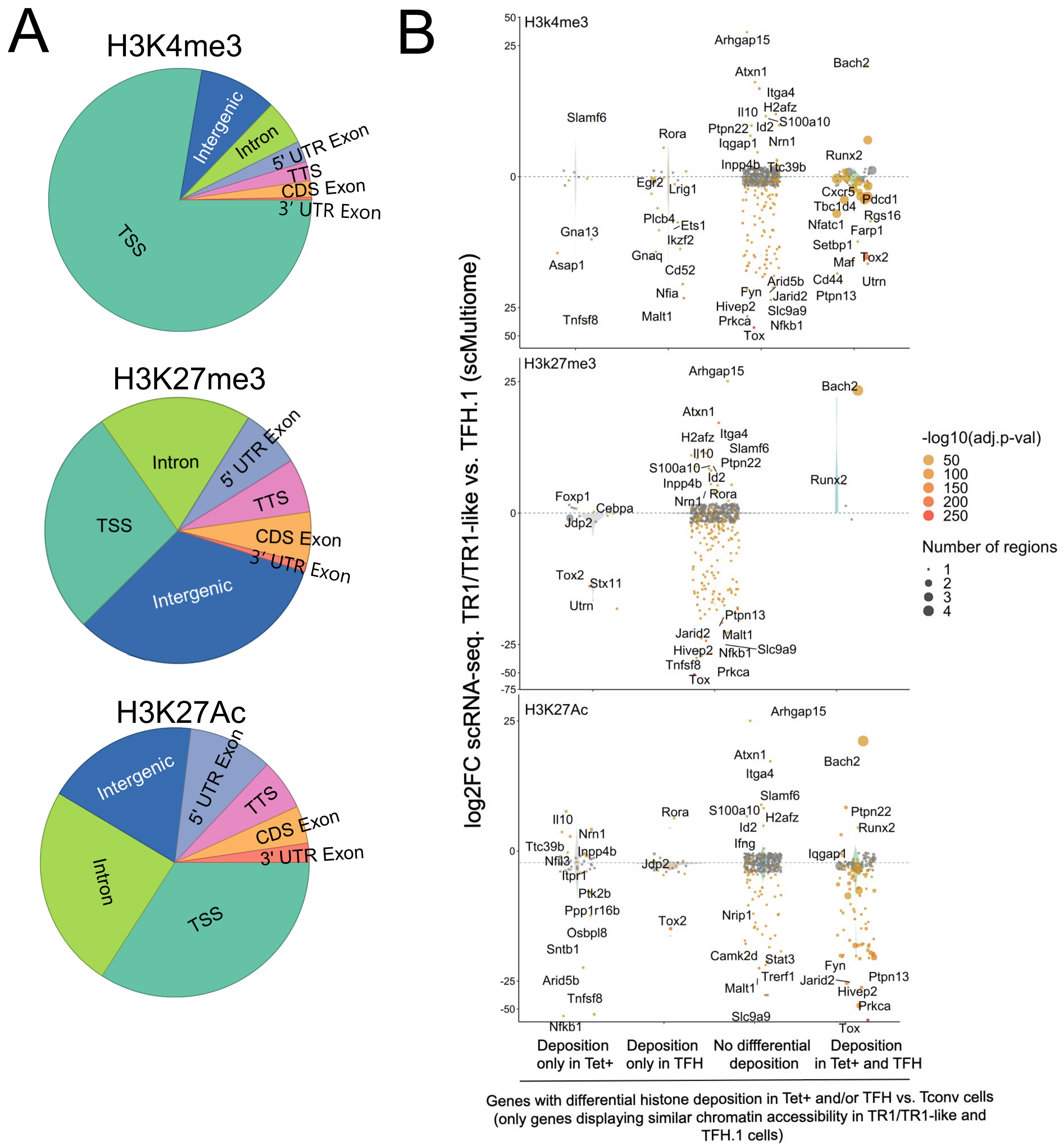
A
H3K4me3
TSS
Intergenic
Intron
5' UTR Exon
TTS
CDS Exon
3' UTR Exon
H3K27me3
TSS
Intron
5' UTR Exon
TTS
CDS Exon
3' UTR Exon
Intergenic
H3K27Ac
Intergenic
5' UTR Exon
TTS
CDS Exon
3' UTR Exon
Intron
TSS
B
H3k4me3
Arhgap15
Bach2
Atxn1
Itga4
H2afz
Il10
Id2
S100a10
Ptpn22
Slamf6
Rora
Iqgap1
Nrn1
Runx2
Inpp4b
Ttc39b
Egr2
Lrig1
Cxcr5
Pdcd1
Tbc1d4
Rgs16
Plcb4
Ets1
Nfatc1
Farp1
Gna13
Ikzf2
Setbp1
Gnaq
Maf
Tox2
Asap1
Cd52
Arid5b
Cd44
Utrn
Nfia
Fyn
Jarid2
Ptpn13
Hivep2
Slc9a9
Malt1
Tnfsf8
Prkca
Nfkb1
Tox
H3k27me3
Arhgap15
Bach2
Atxn1
H2afz
Itga4
Il10
Slamf6
S100a10
Ptpn22
Id2
Runx2
Inpp4b
Foxp1
Rora
Nrn1
Cebpa
Jdp2
Tox2
Stx11
Utrn
Ptpn13
Jarid2
Malt1
Hivep2
Nfkb1
Slc9a9
Tnfsf8
Prkca
Tox
H3K27Ac
Arhgap15
Atxn1
Bach2
Itga4
Slamf6
Il10
Rora
S100a10
H2afz
Ptpn22
Nrn1
Id2
Runx2
Ttc39b
Ifng
Inpp4b
Nfil3
Jdp2
Iqgap1
Itpr1
Ptk2b
Ppp1r16b
Tox2
Nrip1
Osbpl8
Sntb1
Camk2d
Stat3
Fyn
Trerf1
Ptpn13
Arid5b
Jarid2
Malt1
Hivep2
Tnfsf8
Prkca
Slc9a9
Nfkb1
Tox
log2FC scRNA-seq. TR1/TR1-like vs. TFH.1 (scMultiome)
-log10(adj.p-val)
50
100
150
200
250
Number of regions
1
2
3
4
Deposition only in Tet+
Deposition only in TFH
No diffferential deposition
Deposition in Tet+ and TFH
Genes with differential histone deposition in Tet+ and/or TFH vs. Tconv cells (only genes displaying similar chromatin accessibility in TR1/TR1-like and TFH.1 cells)

**Supplementary Fig. 2**. **Differential gene deposition of H3K4me3, H3K27me3, and H3K27ac marks in BDC2.5mi/I-Ag7-NP-induced Tet+ cells, KLH-DNP-induced TFH cells and Tconv cells versus differential gene expression.** (**A**) Pie charts showing the relative distribution of H3K4me3 (top), H3K27me3 (middle), and H3K27Ac (bottom) marks in different gene regions. (**B**) Jitter plots comparing differences in gene expression between BDC2.5mi/I-$A^{g7}$-NP-induced TR1/TR1-like and TFH.1 cells (log2 fold change between the two cell clusters in the scMultiome dataset) for genes linked to OCRs shared between BDC2.5mi/I-$A^{g7}$-NP-induced Tet+ and KLH-DNP-induced TFH cells, as a function of shared or differential H3K4me3 (top), H3K27me3 (middle) and H3K27Ac (bottom) deposition status (FDR<0.01). Gene expression was not associated with differential histone deposition in any case (Wilcox test for differential gene expression: adjusted $P < 0.05$): H3K4me3: Pearson's Chi-square test $P = 0.53$; H3K27me3: Fisher exact test $P=1$; H3K27Ac: Pearson's Chi-square test $P=0.23$. Color depicts the -log10(adjusted p-value) of scRNAseq analysis. Bubble size corresponds to the number of marked regions annotated to specific genes. Most differentially expressed and TR1/TR1-like and TFH-relevant genes included in the 106 genes list from **Supplementary Table 1** are labeled.

Suppl. Fig. 3

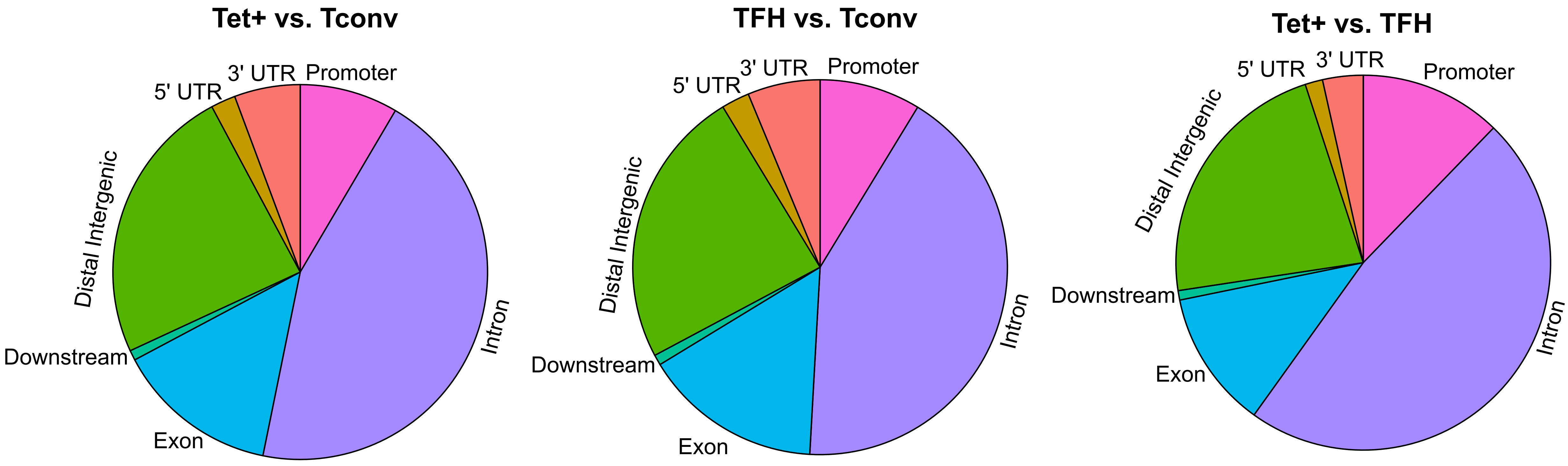
Tet+ vs. Tconv
3' UTR
Promoter
5' UTR
Distal Intergenic
Intron
Downstream
Exon
TFH vs. Tconv
3' UTR
Promoter
5' UTR
Distal Intergenic
Intron
Downstream
Exon
Tet+ vs. TFH
5' UTR
3' UTR
Promoter
Distal Intergenic
Downstream
Intron
Exon

**Supplementary Figure 3**. **Differences in gene region distribution of DMRs among BDC2.5mi/I-A$^{g7}$-NP-induced Tet+, KLH-DNP-induced TFH and Tconv cells.** Pie charts show the relative distribution of DMRs in BDC2.5mi/I-A$^{g7}$-NP-induced Tet+ vs Tconv (left), KLH-DNP-induced TFH vs Tconv (middle), and BDC2.5mi/I-A$^{g7}$-NP-induced Tet+ vs KLH-DNP-induced TFH (right).

Suppl. Fig. 4

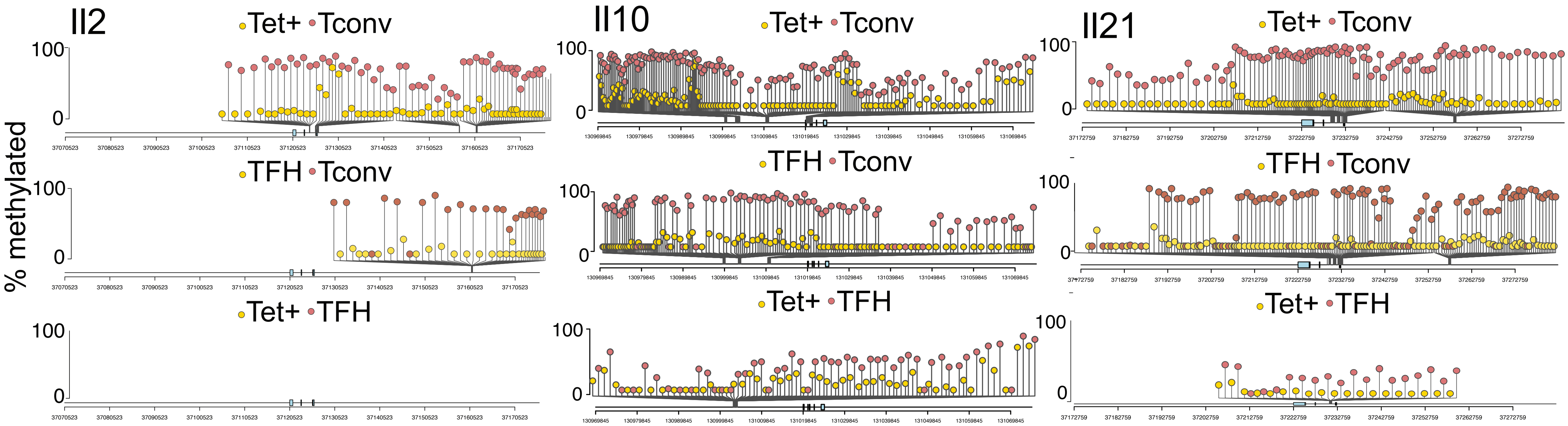

**Supplemental Figure 4. Differential DNA methylation at or near the *Il2*, *Il10* and *Il21* loci between BDC2.5mi/I-A$^{g7}$-NP-induced Tet+ or KLH-DNP-induced TFH and Tconv cells** Lolliplots comparing the location of differentially methylated CpGs in *Il2* (left), *Il10* (middle), and *Il21* (right) in BDC2.5mi/I-A$^{g7}$-NP-induced Tet+ vs Tconv (top), KLH-DNP-induced TFH vs Tconv (middle) and BDC2.5mi/I-A$^{g7}$-NP-induced Tet+ vs KLH-DNP-induced TFH cells comparisons (bottom).

Suppl. Fig. 5

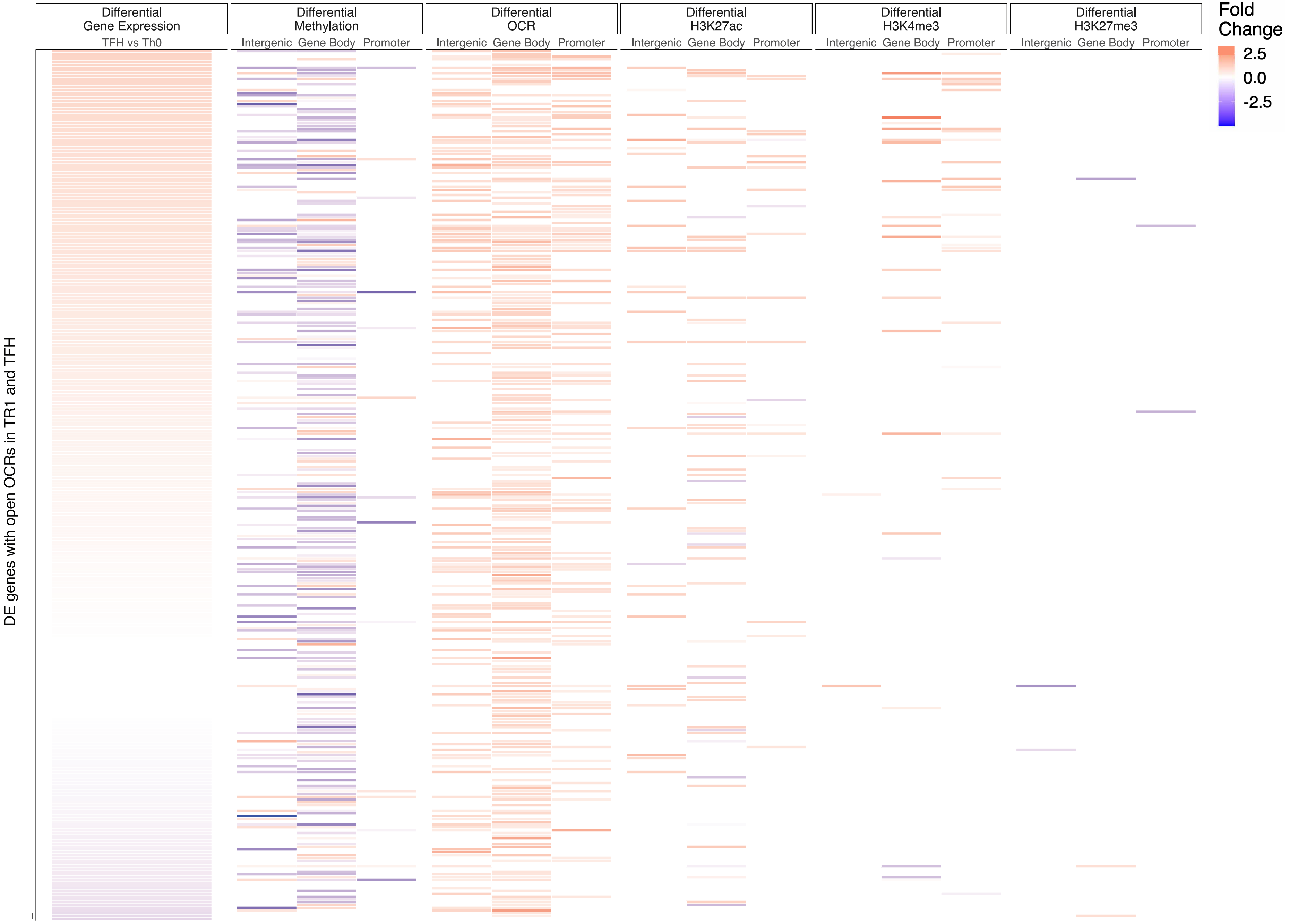

Differential Gene Expression
Differential Methylation
Differential OCR
Differential H3K27ac
Differential H3K4me3
Differential H3K27me3
TFH vs Th0
Intergenic
Gene Body
Promoter
Fold Change
2.5
0.0
-2.5
DE genes with open OCRs in TR1 and TFH

**Supplementary Figure 5**. **Relative contribution of different epigenetic marks to changes in gene expression during the TFH to TR1 cell conversion (TFH vs. Tconv).** Heatmap depicting the presence of different epigenetic marks (from left to right: differential methylation, differential OCRs, differential H3K27ac deposition, differential H3K4me3 deposition and differential H3K27me3 deposition) in antigen induced TFH vs Tconv cells. Data correspond to differentially expressed genes with shared OCRs between TR1 and TFH cells. Differential epigenetic data is scaled for each technique and when multiple genomic regions are associated to a gene, the average is provided. Genes are ordered by decreasing Fold Change of differential gene expression.

Suppl. Fig. 6

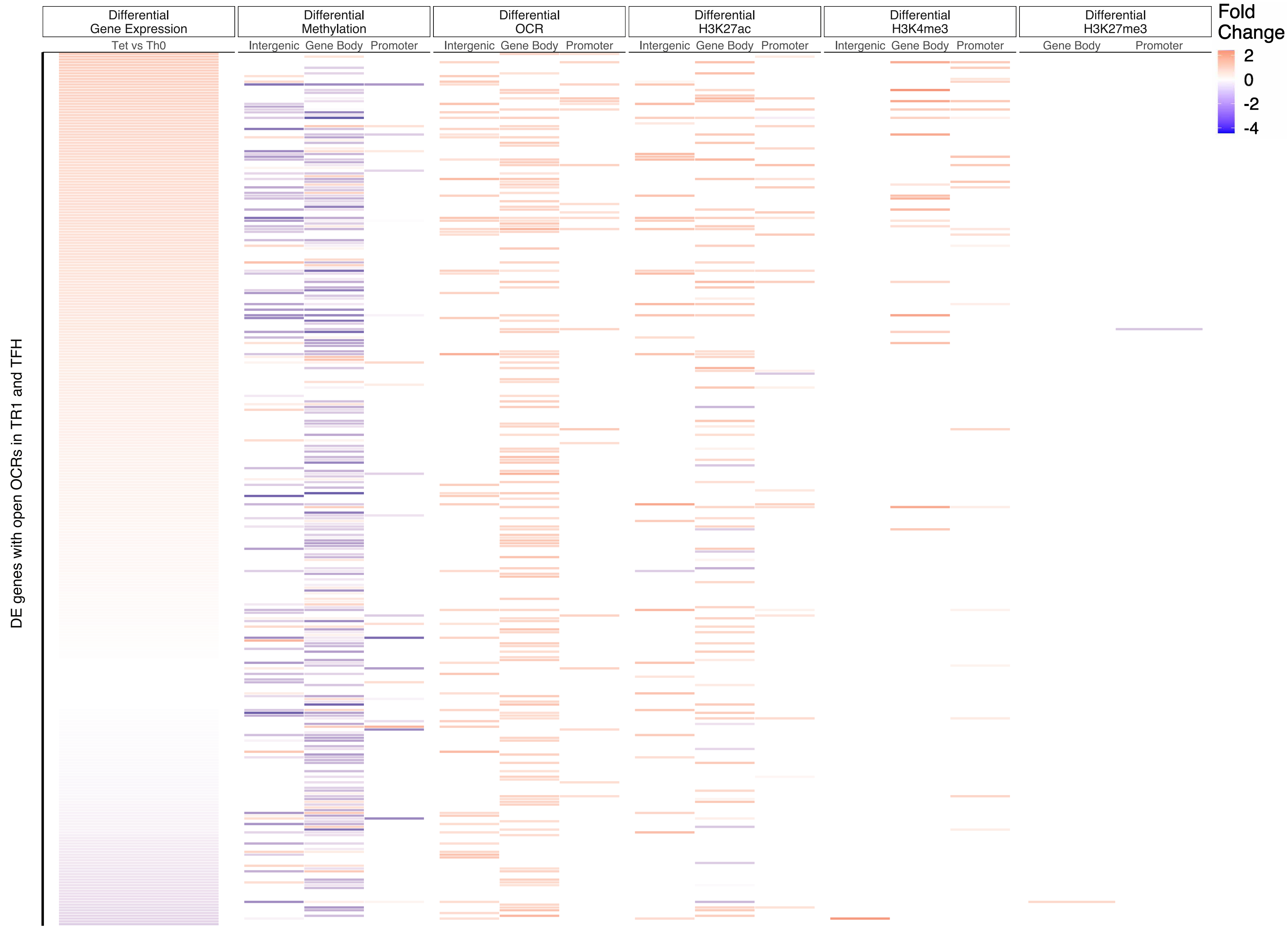

Differential Gene Expression
Tet vs Th0
Differential Methylation
Intergenic Gene Body Promoter
Differential OCR
Intergenic Gene Body Promoter
Differential H3K27ac
Intergenic Gene Body Promoter
Differential H3K4me3
Intergenic Gene Body Promoter
Differential H3K27me3
Gene Body Promoter
Fold Change
2
0
-2
-4
DE genes with open OCRs in TR1 and TFH

**Supplementary Figure 6**. **Relative contribution of different epigenetic marks to changes in gene expression during the TFH to TR1 cell conversion (Tet+ vs Tconv).** Heatmap depicting the presence of different epigenetic marks (from left to right: differential methylation, differential OCRs, differential H3K27ac deposition, differential H3K4me3 deposition and differential H3K27me3 deposition) in BDC2.5mi/IA$^{g7}$-NP-induced Tet+ vs Tconv cells. Data correspond to differentially expressed genes with shared OCRs between TR1 and TFH cells. Differential epigenetic data is scaled for each technique and when multiple genomic regions are associated to a gene, the average is provided. Genes are ordered by decreasing Fold Change of differential gene expression.

Suppl. Fig. 7

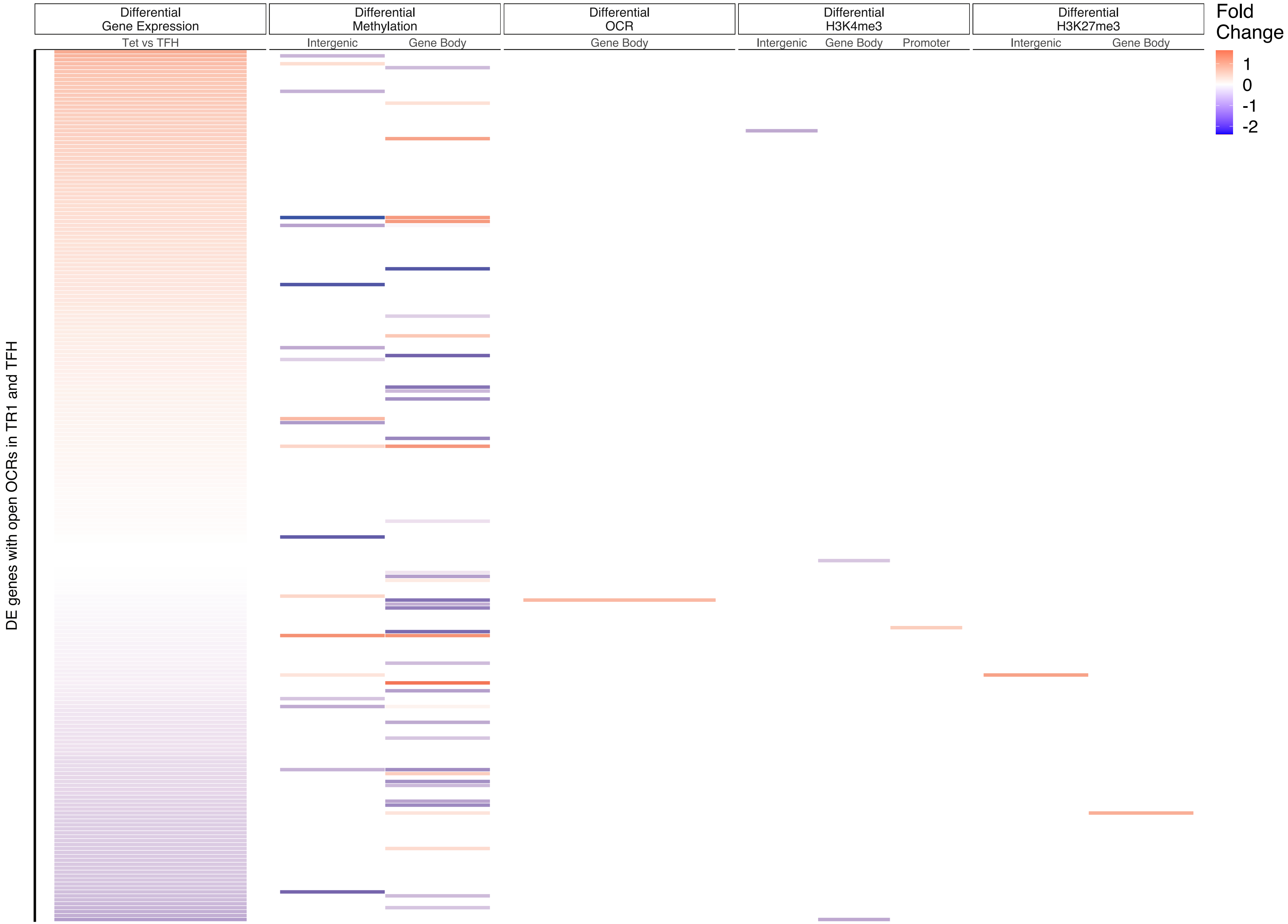
Differential Gene Expression
Tet vs TFH
Differential Methylation
Intergenic
Gene Body
Differential OCR
Gene Body
Differential H3K4me3
Intergenic
Gene Body
Promoter
Differential H3K27me3
Intergenic
Gene Body
Fold Change
1
0
-1
-2
DE genes with open OCRs in TR1 and TFH

**Supplementary Figure 7**. **Relative contribution of different epigenetic marks to changes in gene expression during the TFH to TR1 cell conversion (Tet+ vs TFH).** Heatmap depicting the presence of different epigenetic marks (from left to right: differential methylation, differential OCRs, differential H3K27ac deposition, differential H3K4me3 deposition and differential H3K27me3 deposition) in in BDC2.5mi/IA$^{g7}$-NP-induced Tet+ vs KLH-DNP-induced TFH cells. Data correspond to differentially expressed genes with shared OCRs between TR1 and TFH cells. Differential epigenetic data is scaled for each technique and when multiple genomic regions are associated to a gene, the average is provided. Genes are ordered by decreasing Fold Change of differential gene expression.
